# Alternating dynamics of *oriC*, SMC and MksBEF in segregation of *Pseudomonas aeruginosa* chromosome

**DOI:** 10.1101/2020.05.23.112730

**Authors:** Hang Zhao, Bijit Bhowmik, Valentin V. Rybenkov

## Abstract

Condensins are essential for global chromosome organization in diverse bacteria. Atypically, *Pseudomonas aeruginosa* encodes condensins from two superfamilies, SMC-ScpAB and MksBEF. We report that the two proteins play specialized roles in chromosome packing and segregation and are synthetically lethal with ParB. Inactivation of SMC or MksB asymmetrically affected global chromosome layout, its timing of segregation and sometimes triggered a chromosomal inversion. Localization pattern was also unique to each protein. SMC clusters colocalized with *oriC* throughout cell cycle except shortly after origin duplication, whereas MksB clusters emerged at cell quarters shortly prior to *oriC* duplication and stayed there even after cell division. Relocation of the proteins was abrupt and coordinated with *oriC* dynamic. These data reveal that the two condensins asymmetrically play dual roles in chromosome maintenance by organizing it and mediating its segregation. Furthermore, the choreography of condensins and *oriC* relocations suggest an elegant mechanism for the birth and maturation of chromosomes.

**Importance:** Mechanisms that define the chromosome as a structural entity remain unknown. A key element in this process are condensins, which globally organize chromosomes and contribute to their segregation. This study characterized condensin and chromosome dynamics in *Pseudomonas aeruginosa*, which harbors condensins from two major protein superfamilies, SMC and MksBEF. The study revealed that both proteins asymmetrically play a dual role in chromosome maintenance by spatially organizing the chromosomes and guiding their segregation but can substitute for each other in some activities. The timing of chromosome, SMC and MksBEF relocation was highly ordered and interdependent revealing causative relationships in the process. Moreover, MksBEF was found to produce clusters at the site of chromosome replication that survived cell division and remained in place until chromosome replication was complete. Overall, these data delineate the functions of condensins from the SMC MksBEF superfamilies, reveal the existence of a chromosome organizing center and suggest a mechanism that might explain the biogenesis of chromosomes.

---

*Pseudomonas aeruginosa* is a significant opportunistic human pathogen responsible for a variety of infectious diseases in immunocompromised patients and those with breached cutaneous barrier (1, 2). The high pathogenicity of *P. aeruginosa* resides in its numerous encoded virulence factors, a cell envelope that is virtually impermeable to most available antibiotics, a plethora of metabolic enzymes suitable for diverse niches, and a sophisticated signaling network that accelerates adaptation of the bacterium to hostile environment (3–6). Recent studies revealed that the global chromosome dynamics in *P. aeruginosa* is also under an elaborate control that is integrated into epigenetic and virulent behavior of the bacterium (7).

A key element in this system are condensins, the characteristically V-shaped ABC type ATPases that form macromolecular clamps on DNA that can bridge distant DNA fragments and have an intrinsic ability to assemble into a macromolecular scaffold at the core of chromosomes (reviewed in (8–10)). In bacteria, these proteins form dynamic clusters in the middle of nucleoids where they presumably act as chromosome organizing centers (11–13). Condensins act in cooperation with ubiquitous other nucleoid associated proteins to impose a multilayer organization of the bacterial chromosome (14, 15).

Three families of condensins have been identified in bacteria. The oldest known condensin, MukBEF, is found in *E. coli* and related γ-Proteobacteria (16). Most other bacteria carry the SMC-ScpAB complex (17). Its SMC subunit is highly homologous to the archaeal and eukaryotic proteins, but not MukB. Members of the third family, MksBEF, sporadically occur in diverse bacteria, including many environmental and pathogenic strains, typically in combination with SMC-ScpAB or MukBEF (18). Several families MksBEF were discovered, each with a distant homology to another and some traceable to the *E. coli* MukBEF. In this respect, MukBEF and MksBEF can be viewed as a large and diverse superfamily of condensins.

Despite sequence divergence and a number of biochemical distinctions, MukBEF and SMC appear to play similar roles in their host bacteria. Both proteins are able to bridge DNAs, form clusters at the core of the chromosome, which relocate within the cell in parallel with duplicating chromosomes, and are essential for faithful chromosome segregation (9, 19, 20). However, their DNA recruitment mechanism appears idiosyncratic to the protein. SMC proteins are loaded onto the chromosome in the vicinity of the origin or replication, *oriC*, with the help of the ParABS system (21, 22), whereas MukBEF employs DNA topoisomerase IV and MatP for its recruitment to the *ori* and *ter* domains of *E. coli*, respectively (23–25).

To highlight the differences between the two condensins, we explored chromosome segregation in *P. aeruginosa*, which encodes both SMC and MksBEF and where the defects of condensin inactivation are reasonably well tolerated. The former factor helps reveal the differences in protein localization, if any, whereas the latter one reduces the secondary effects of chromosome disorganization caused by condensin inactivation. These benefits of the system allow one to address a cornerstone question in condensin biology, whether the proteins provide the driving force for chromosome segregation or simply follow the segregating chromosomes.

The chromosome of *P. aeruginosa* is longitudinally organized with the origin of replication, *oriC*, located at 80% of the cell length and the replication terminal region at the opposite, newly formed pole of the cell. Replication occurs in the middle of the cell, where the replisome remains for the duration of the cell cycle (26, 27). Segregation begins at the *oriC* locus and proceeds bidirectionally along the two chromosomal arms. Unlike *E. coli*, *P. aeruginosa* lacks the *ter*-Tus system (8). Instead, segregation ends in the vicinity of *dif*, the site of the chromosome dimer resolution. This arrangement is maintained even in the asymmetric chromosome of *P. aeruginosa* strain PAO1-UW where the two chromosomal arms differ by 60%. In contrast, replication proceeds symmetrically along the two arms (27).

We show here that this pattern of segregation is asymmetrically dependent on the presence of condensins, and the effect differs between the *oriC* region and the rest of the chromosome. In particular, the bulk chromosome segregation was delayed in MksB-deficient cells but occurred sooner in a *smc* mutant. For *oriC*, however, the deletion of any of the condensins produced the same effect, a delay in segregation of one of the sister origins of replication. The deletion of *smc* or *mksB* dispersed the large domains previously found in the longer arm of the PAO1-UW chromosome. Moreover, MksB-deficient cells underwent a large chromosomal inversion, which equalized the length of its arms and realigned *dif* and the terminus of replication. In *smc* mutants, segregation proceeded symmetrically and terminated opposite from *oriC* rather than at *dif*. Thus, SMC and MksB have asymmetric functions in global chromosome folding and segregation. Accordingly, MksB and SMC clusters rarely colocalized with each other. While SMC was primarily found at *oriC*, MksB was located at the midcell for most of the cell cycle. Relocation of both proteins was tied to *oriC* segregation suggestive of a cause-and-effect relationship in the process. Overall, these data suggest that condensins comprise the core of a chromosome organizing center in *P. aeruginosa*, which begins its life at a newly formed origin of replication and later matures into an assembly in the middle of the cell.

## Results

### SMC and MksBEF form separate clusters within the chromosome

We first examined subcellular localization of GFP-tagged MksB and SMC in *P. aeruginosa* cells grown in rich medium. Both proteins formed distinct clusters in the middle of short cells and about quarter positions of longer cells (Fig. 1A). Such localization is characteristic of bacterial condensins (11–13) and is also reminiscent of *oriC*-proximal regions of the *P. aeruginosa* chromosome (26). Both MksB and SMC were found in the middle of nucleoids (Fig. 1D), which is further consistent with their expected role in chromosome organization. Notably, MksB foci duplicated later than SMC foci (Fig. 1B, C), indicating that they associate with different intracellular targets and might be playing distinct roles.

**Figure 1.**
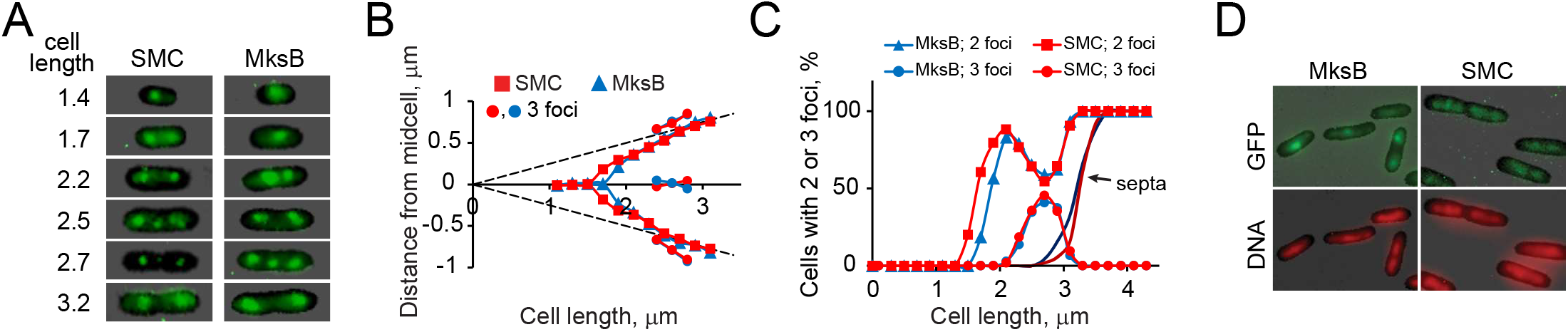
Subcellular localization of MksB and SMC in rapidly growing *P. aeruginosa*. (**A**) Focal localization of GFP-tagged MksB and SMC. Cell length is measured in μm. (**B**). Average position of MksB and SMC foci in cells (*n* > 500) with two or three foci. Localization of the foci was quantified using Nucleus program (38). (**C**) Predominance of cells with two or three foci. Also shown are the proportions of cells containing a septum. Note that the formation of the three-focus cells parallels a decline in two-focus cells (**D**). Colocalization of MksB-GFP and SMC-GFP with DNA.

Shortly before septum constriction, a third focus could be observed in the middle of the cell. This focus formed only transiently, after the split of the main foci, and soon dispersed at the onset of septum constriction (Fig. 1C). MksB and SMC produced the third focus simultaneously, suggesting the same mechanism for its formation. In all cases, the third focus was found in the middle of a DNA mass and not at a periphery of the nucleoids (Fig. 1D). Moreover, its location in the middle of the cell suggested other attractors than duplicating *oriC*s in the daughter cells.

Piqued by these unusual features, we further explored chromosome segregation in *P. aeruginosa*.

### Deletion of condensins desynchronizes segregation of *oriC*

Subsequent experiments were carried out with cells grown in minimal media, when replication is initiated only once per cell cycle. *OriC*-proximal region was tagged with *tetO* repeats and visualized using fluorescent repressor operator system, FROS, as previously described (26, 27). Its location was followed in parental PAO1-UW cells and cells deficient in one or both condensins. As previously shown (26, 27), the chromosome in *P. aeruginosa* is stretched along the major axis of the cell from *oriC* to *dif*. At the onset of replication, *oriC* moves to the midcell, duplicates, the daughter *oriC*’s then move apart, and the mother chromosome is gradually pulled towards the replisome located in the middle of the cell (Fig. 2A).

**Figure 2.**
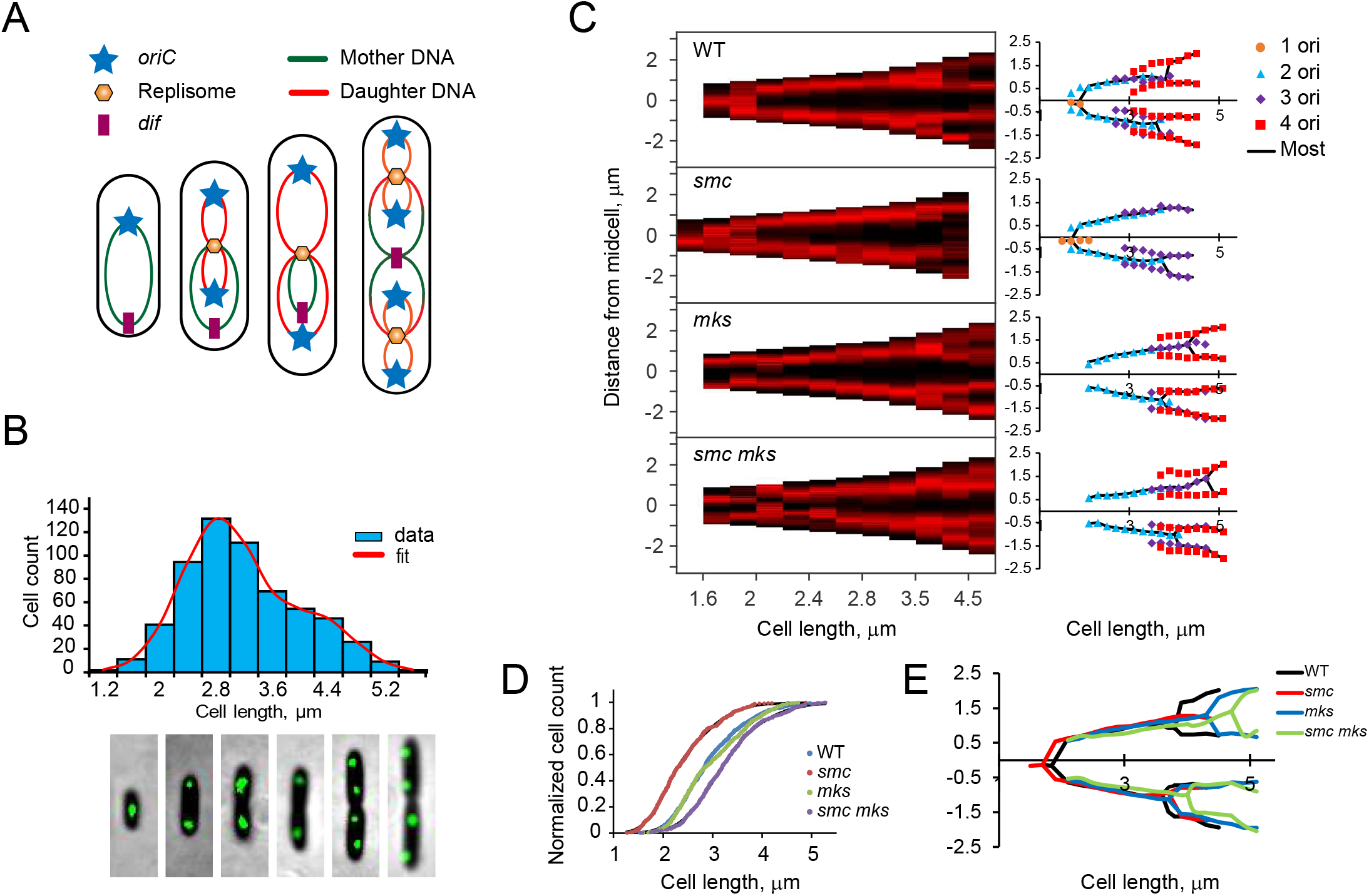
Segregation of the origin of replication in condensin deficient cells. (**A**) Layout of *P. aeruginosa* chromosome during replication. (**B**) Location of *oriC* in representative cells (bottom) matched with the bimodal distribution of cell sizes (top). (**C**) Binned demographs of condensin variants (left, *n*>300) and average position of *oriC* as a function of cell length (right). On demographs, the cell outline is shown in black and the intensity of *oriC* fluorescence in red. Black line on the averages graphs traces position of *oriC* in the dominant population of cells. (**D**) Cumulative cell size distribution in wild type and condensin deficient cells. Data were fit to an integrated double Gaussian distribution. (**E**) Comparison of *oriC* segregation in condensin- proficient and deficient cells.

In agreement with previous studies, most of the newly born *P. aeruginosa* cells contained a single *oriC* (Fig. 2B, C). When cells grew up to 1.6 m, *oriC* relocated to the middle of the cell. Duplication of *oriC* occurred at cell length of 1.8 m, after which sister *oriC’s* moved apart and then gradually migrated towards the 20%/80% positions. At cell length of 3.6 m, the second round of *oriC* duplication could be observed. Cell division did not correlate with *oriC* duplication and could occur before or after it, giving birth to shorter cells with a single *oriC* or longer ones with two origins of replication. A delay in cell division relative to the second round of chromosome replication probably explains the bimodal distribution of cell lengths (Fig. 2B).

Deletion of *smc* altered the aforementioned pattern in two ways. First, the average cell length declined to 2.3 m, compared to 2.8 m found in wild type cells (Fig. 2D). Second, segregation of only one of the sister *oriC*’s could be observed prior to cell division, resulting in a large number of newborn cells with a single origin of replication (Fig. 2C, second panel).

Deletion of *mksB* had an opposite effect on cell morphology. Although the average cell length was similar to that in the parental strain (Fig. 2D), the cell division event was delayed relative to the second round of origin duplication (Fig. 2C, third panel). As a result, virtually no newborn cells with a single *oriC* could be found. Moreover, we observed a pronounced delay in separation of the two sister *oriC’s*. Although not entirely synchronous, the two *oriC*’s in wild type cells separated over a narrow range of cell lengths, averaging 3.6 and 3.8 m for the first and second event, respectively. In *mksB* cells, the lengths were 3.9 and 4.5 m.

Remarkably, the deletion of both condensins did not produce dramatic effects on *oriC* segregation. The morphology of *smc mksB* cells was similar to that of the *mksB* mutant, except that a delay in segregation of *oriC*’s became more pronounced (Fig. 2C, E) and the average cell length increased to 3.2 m (Fig. 2D). Thus, none of the condensins is required for segregation of *oriC*’s, alone or together; however, both are needed to ensure the synchronicity of *oriC* segregation in the two daughter cells.

### A delay between replication and segregation of *oriC*’s

The found delay in chromosome segregation could in principle be caused by a delay in replication. To evaluate this possibility, we explored chromosome segregation in condensin mutants using marker frequency analysis. To this end, chromosomal DNA of the mutant cells was subjected to deep sequencing and the relative abundances of genomic loci fit to a model that postulates up to two replication start events at *oriC* during the life time of the cell (27). This analysis quantifies duplication of genomic loci following the passage of a replication fork, which leads to a higher copy number of origin-proximal DNA fragments (Fig. 3A).

**Fig. 3.**
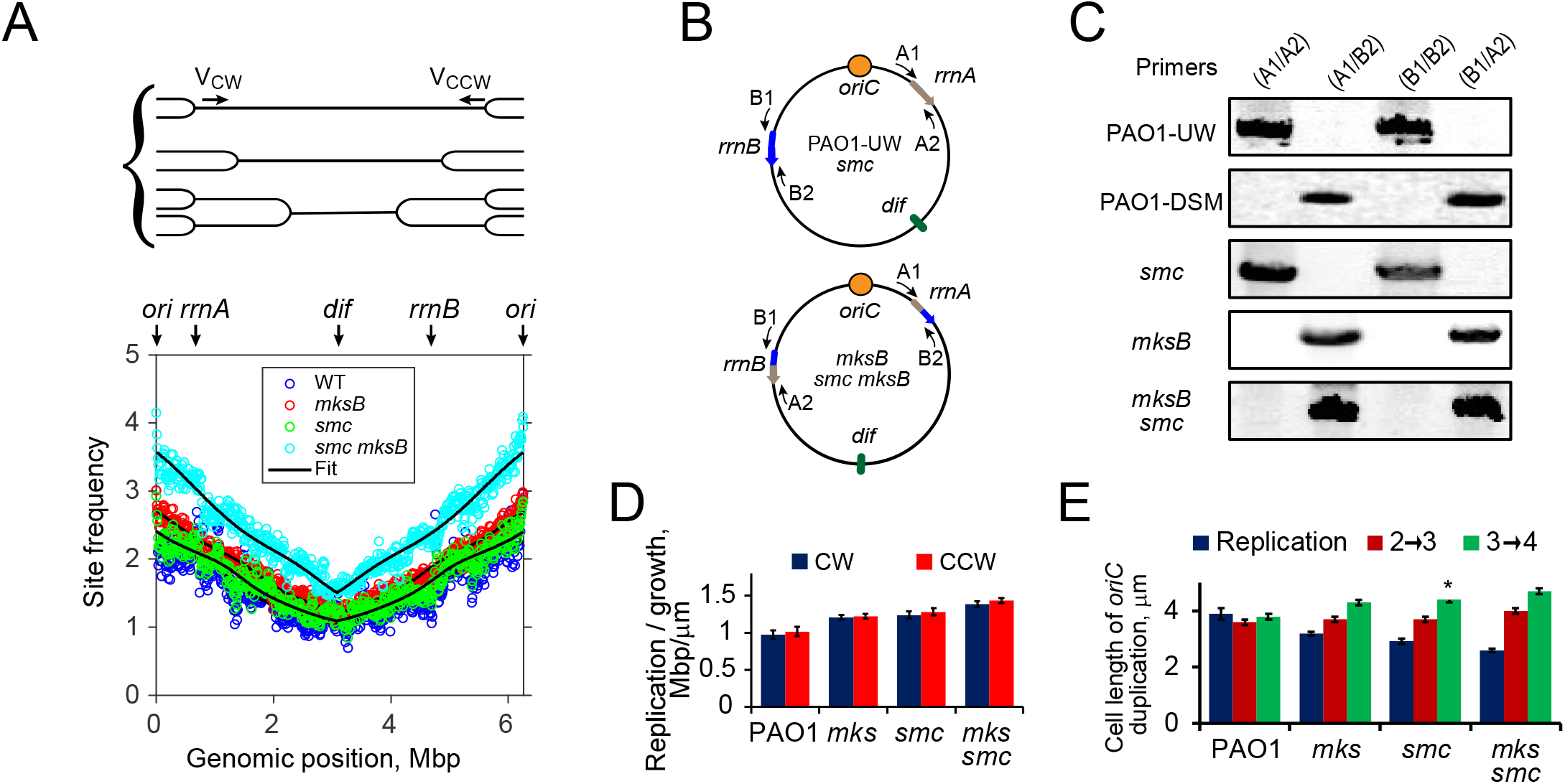
Chromosome replication in the presence and absence of condensins. (**A**) Marker frequency analysis of chromosome replication. Cell culture contains multiple cell populations where bidirectional DNA replication advanced to its own extent, resulting in a higher copy number of origin-proximal regions (top). Bottom panel shows frequencies of all detected chromosomal sites for the four condensin variant strains fit to the model that postulates a constant rate of replication. For *mksB* and *smc mksB* strains, the genomic position takes into account the inversion between *rrnA* and *rrnB*. (**B**, **C**) PCR analysis of the genomic connectivity at *rrnA* and *rrnB* sites. Primers A1, A2, B1 and B2 are homologous to genomic regions flanking the rDNA sites (**B**). Both MksB-deficient strains carry the same inversion as found in PAO1-DSM (**C**). (**D**) The best-fit chromosome replication rates for the clockwise (CW) and counterclockwise (CCW) forks normalized to the cell growth rate for the parental strain and condensin mutants. (**E**) The best-fit cell length of *oriC* replication in condensin mutants compared to the cell length at which *oriC*’s are segregated in the two daughter cells. The segregation data are as determined in Fig. 2E for the transitions from two- to three-*oriC* cells (2→3) and from three- to four-*oriC* cells (3→4). For *smc* mutants, no second *oriC* duplication was ever observed, and the bar represents the low bound on the expected cell length at the duplication (*).

Unexpectedly, we found discontinuities in the marker frequency patterns of *mksB* and *smc mksB* cells (Fig S1). PCR analysis of the implicated region revealed that they are caused by a large chromosomal inversion between the *rrnA* and *rrnB* regions, which occurred after the deletion of *mksB* gene (Fig 3B, C). This inversion was found in all 25 analyzed *ΔmksB* colonies obtained after 5 separate conjugations, suggesting that it is tightly linked to *mksB*. Notably, the same inversion was reported for the PAO1-DSM lineage of PAO1 strains, where it is accompanied by a deletion in *mksE*. Apparently, a fully functional MksBEF is required to maintain the inversion found in PAO1-UW cells. The existence of this inversion is accounted for in further analysis.

In all three mutants, replication started at *oriC* and proceeded bidirectionally to terminate opposite from *oriC*, indicating that the clockwise and counter-clockwise forks advance with the same rate (Fig. 3D). However, the rate of replication varied from one mutant to another. The deletion of *smc* or *mksB* resulted in 20% faster replication, relative to cell growth, than for the wild type cells, whereas the effect was additively higher, 40%, for the double mutant (Fig. 3D).

Accordingly, *oriC* replication started earlier in condensin mutants than in the parental strain (Fig. 3E). In wild type cells, the best-fit start of *oriC* replication was simultaneous with segregation of the origin in both daughter cells. In contrast, we observed a clear delay in segregation of *oriC*’s following their replication, which became especially pronounced in the double condensin mutant. Thus, the delay in *oriC* segregation in one of the sister cells (Fig. 2) cannot be attributed to *oriC* replication.

### Inactivation of SMC and MksB dissolves all domains, disorganizes *dif* region

We next introduced fluorescent tags into various locations of *P. aeruginosa* chromosome and examined chromosome segregation in condensin deficient cells. As reported previously for the wild type *P. aeruginosa* (27), chromosomal loci of both *smc* and *mksB* mutants migrated towards the middle of the cell, where they duplicated and moved apart (Fig. 4A, B). Foci duplication occurred sequentially in accord with their genomic location. The second round of segregation was only detected for the *oriC* region. To facilitate the comparison of all tagged chromosomal sites, we next marked the midpoints of segregation on the map of the chromosome and connected the DNA sites on the opposite arms that segregated at the same cell length (Fig. 4C, D). In the parental PAO1-UW strain, chromosome segregation proceeds sequentially from *oriC* to *dif* along the two chromosomal arms, with the exception or two large domains on the longer arm, which segregate simultaneously ((27), Fig. 4E). No such discontinuities were observed in MksB- or SMC-deficient strains (Fig. 4C, D). Thus, all macrodomains previously observed in PAO1-UW cells were dispersed following inactivation of either of the condensins.

**Fig. 4.**
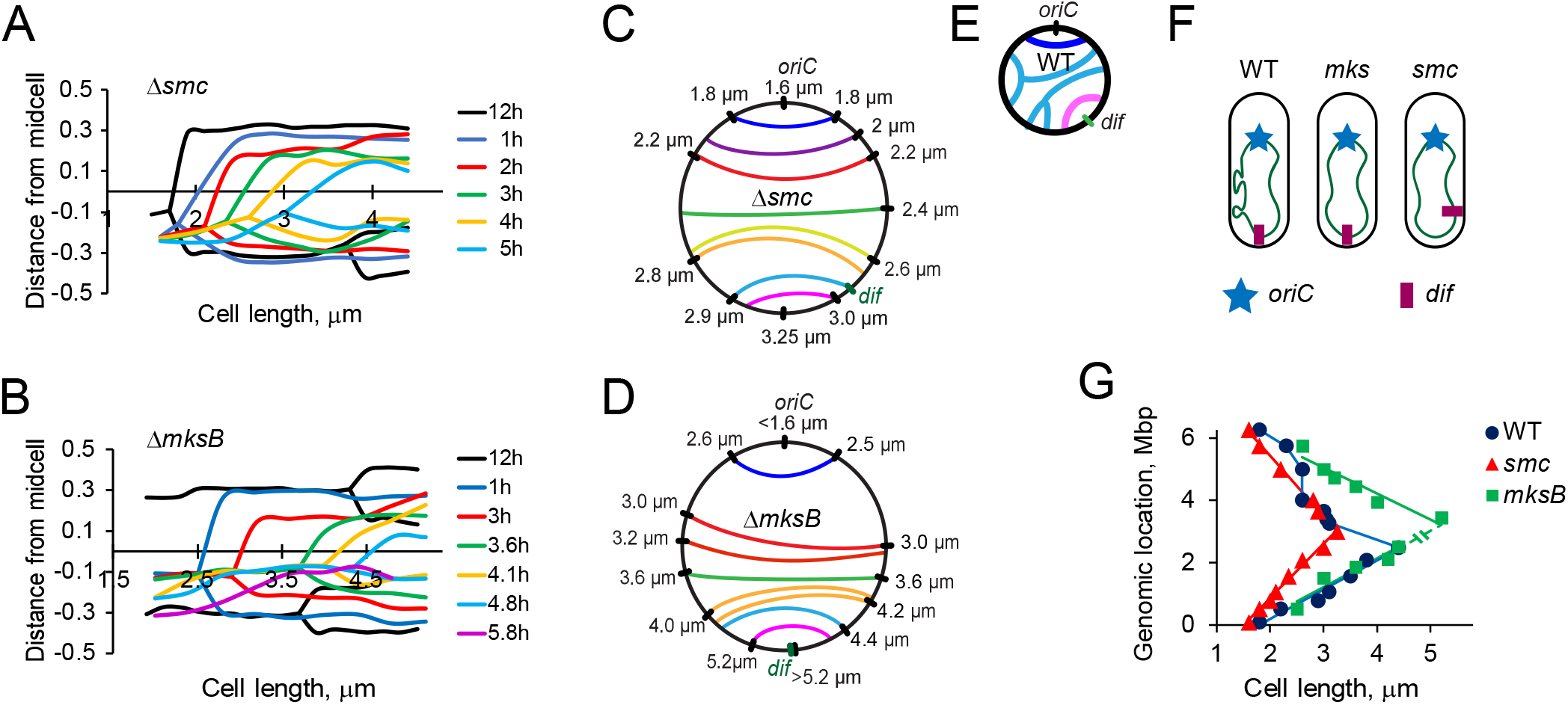
Divergent effects of SMC and MksB inactivation on chromosome segregation. (**A**, **B**) Segregation of the tagged chromosomal loci in SMC- (**A**) or MksB- (**B**) deficient cells. The intracellular positions of each fluorescently tagged locus is shown as a function of cell length. (**C**, **D**) Dial graphs of chromosome segregation which connect sites on the two chromosomal arms that segregate at the same cell length. (**E**) A dial graph for the parental strain highlighting two large domains on the longer arm. (**F**) A diagram summarizing chromosome segregation patterns in wild type and condensin-deficient cells. (**G**) A summary of chromosome segregation in the three strains. Note that segregation of *dif* was never detected in *mksB* cells.

Another striking difference between condensin mutants and parental cells was in the site of segregation termination. In wild type and MksB-deficient cells, the last segregating region was found in the vicinity of *dif* (Fig. 4D). In *mksB* cells, however, this region is opposite from *oriC* owing to the large chromosomal inversion. This region segregated very late during cell division, and we never found *mksB* cells with two *dif* foci (Fig. 4B, F, G). Apparently, the duplicated *dif* sites remain close to the septum up until the cell division event. This contrasts the parental cells, where separation of *dif* foci was routinely observed prior to cell division. We also readily observed segregation of all tagged sites, including *dif*, in SMC-deficient cells (Fig. 4C, G). In this case, however, *dif* was no longer the last segregated site. Thus, inactivation of either SMC and MksB resulted in disorganization of segregation of the *dif* region.

The effect of condensin deletions could also be seen in the rest of the chromosome (Fig. 4G). In MksB deficient cells, chromosome segregation was markedly delayed compared to the wild type, primarily in the left chromosomal arm. In *smc* cells, chromosome segregation happened sooner, in both arms, consistent with the smaller length of these cells (Fig. 2D).

### Alternating dynamics of *oriC*, SMC and MksB

To further contrast the roles of SMC and MksB in chromosome segregation, we examined localization patterns of the proteins relative to *oriC*. Figure 5A shows binned demographs of cells harboring mCherry-tagged *oriC* and SMC-mVenus. Fluorescence intensity was recorded along the major axis of each cell, cells were binned according to their length, and the intensity averaged within each bin. The binned intensity profiles were plotted as a function of cell length as images against black projections of cells (Fig. 5A). Representative doubly labeled cells are also shown in Fig. S2.

**Fig. 5.**
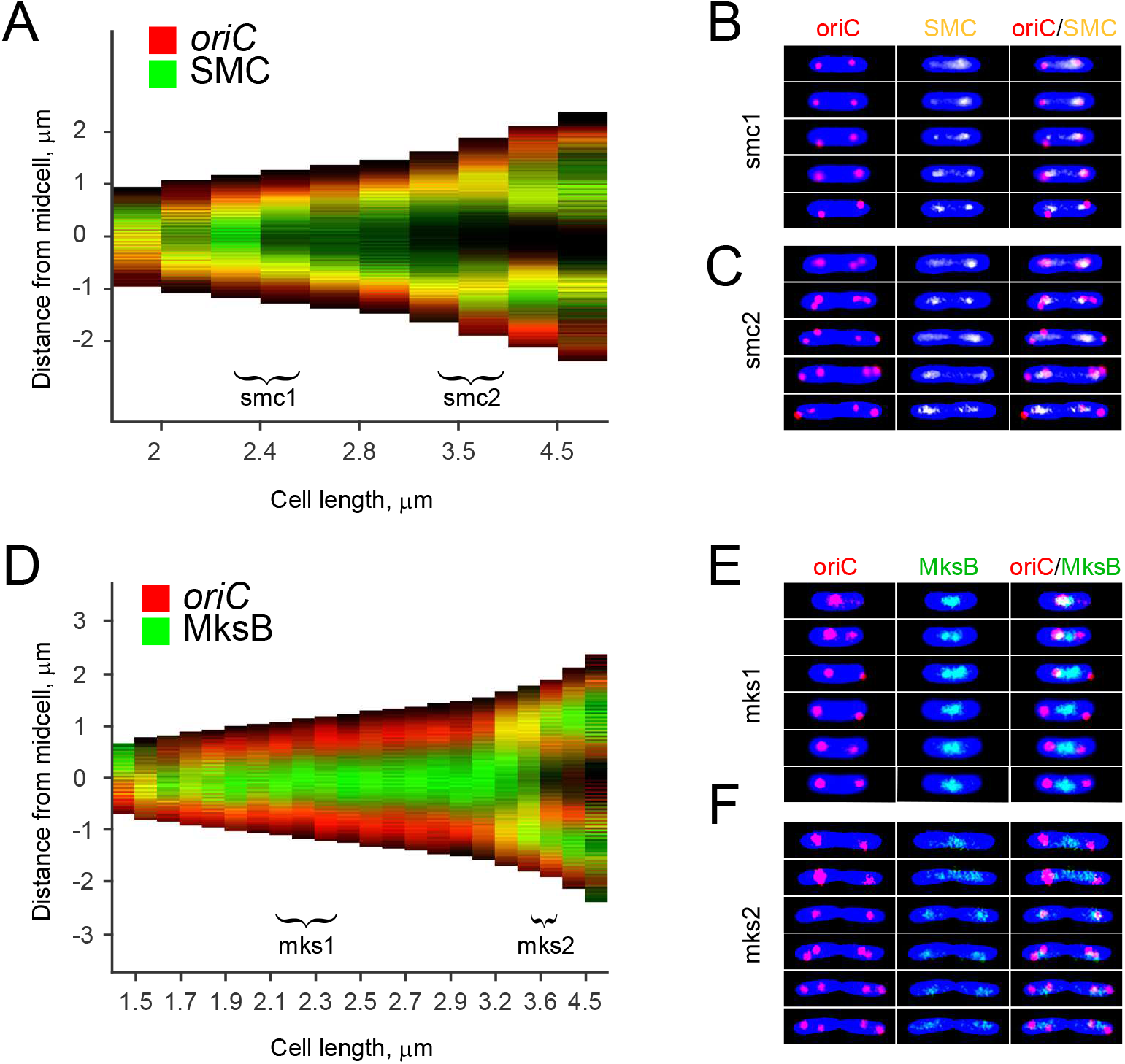
Choreography of *oriC*, SMC and MksB dynamics. (**A**) Binned demographs of cells harboring SMC-mVenus and mCherry-tagged *oriC* (*n*=483). Brackets mark the ranges analyzed in panels B and C. (**B** and **C**) Representative time-lapse images collected every three minutes for cells with initial length of 2.3 μm (**B**) and 3.4 μm (**C**). (**D**) Binned demographs of cells harboring MksB-GFP and mCherry-tagged *oriC* (*n*=1612). (**D** and **E**) Representative time-lapse images collected every three minutes for cells with initial length of 2.1 μm (**E**) and 3.5 μm (**F**).

The shortest cells contained a single *oriC*, which overlapped with a broader single spot of SMC. As the cells grew, *oriC* duplicated, and the two sister *oriC*’s quickly moved apart towards positions at 20% and 80% of the cell length. Most of the SMC-mVenus fluorescence followed one of the *oriC*’s, whereas the other retained only a small fraction of it. Soon thereafter, SMC equalized at the two spots and remained there until the rest of the cell cycle. The sequence of events repeated itself in longer cells, at the next round of *oriC* duplication. The asymmetry of segregation was even clearer in this case. In both daughter chromosomes, the septum-proximal *oriC* retained most of SMC, whereas the poleward origins were slow to recruit the condensin. These data reveal that SMC relocation follows rather than leads relocation of *oriC*. Moreover, the dynamics of SMC appears tied to the overall cell morphology.

The dynamics of MksB was strikingly different. MksB and *oriC* did not colocalize in the shortest newborn cells. They did converge onto the midcell as the cells started growing. Once *oriC* duplicated and moved away, however, it also departed from MksB, which remained in the middle of the cell for the duration of the cell cycle. Curiously, the MksB cluster became unstable shortly before the next round of *oriC* duplication. A new MksB spot appeared at one of the *oriC*’s to be followed soon by *oriC* segregation. Segregation of the second *oriC* occurred in a similar manner, with MksB cluster emerging at the quarter position shortly prior to *oriC* duplication.

Other than that, there was no particular order to the events in the sister half-cells, which appeared to occur independent of each other. Once MksB clusters formed at both cell quarters, the one at the midcell faded away. The subsequent cell division completed the cycle by producing daughter cells with a single MksB cluster located at or close to the middle of the cell.

### Condensins are synthetically lethal with ParB

The prominent role of SMC and MksB in chromosome organization appears at odds with the modest morphological changes caused by their deletion. To address this apparent incongruity, we examined genetic interactions of condensins with the ParAB*S* system. We readily constructed strains deficient in ParB and any one of the condensins but could not construct the triple deletion strain. To confirm that this was indeed due to synthetic lethality of ParB, SMC and MksB, we employed two separate approaches.

We first constructed a merodiploid *ΔparB Δsmc mksB::pEX18-sacB-ΔmksB-Gm* strain, which contains a functional *mksB*. Plating these cells on sucrose selects for recombinants with an excised plasmid backbone. The excision could proceed forward, completing the deletion of *mksB* and rendering cells resistant to gentamicin, or backwards, restoring the endogenous *mksB* gene and generating gentamicin sensitive strains (Fig. 6A). In principle, the probabilities of the two events should be equal, as we found with an unrelated operon MexGHID (Fig. 6B). With MksB, however, all recombination events restored the *mksB^+^* genotype (Fig. 6B). The same result was observed for a merodiploid *ΔparB ΔmksB smc::pEX18-sacB-Δsmc-Gm*, when we tried to remove the *smc* gene from *ΔparB ΔmksB* cells (Fig. 6B).

**Figure 6.**
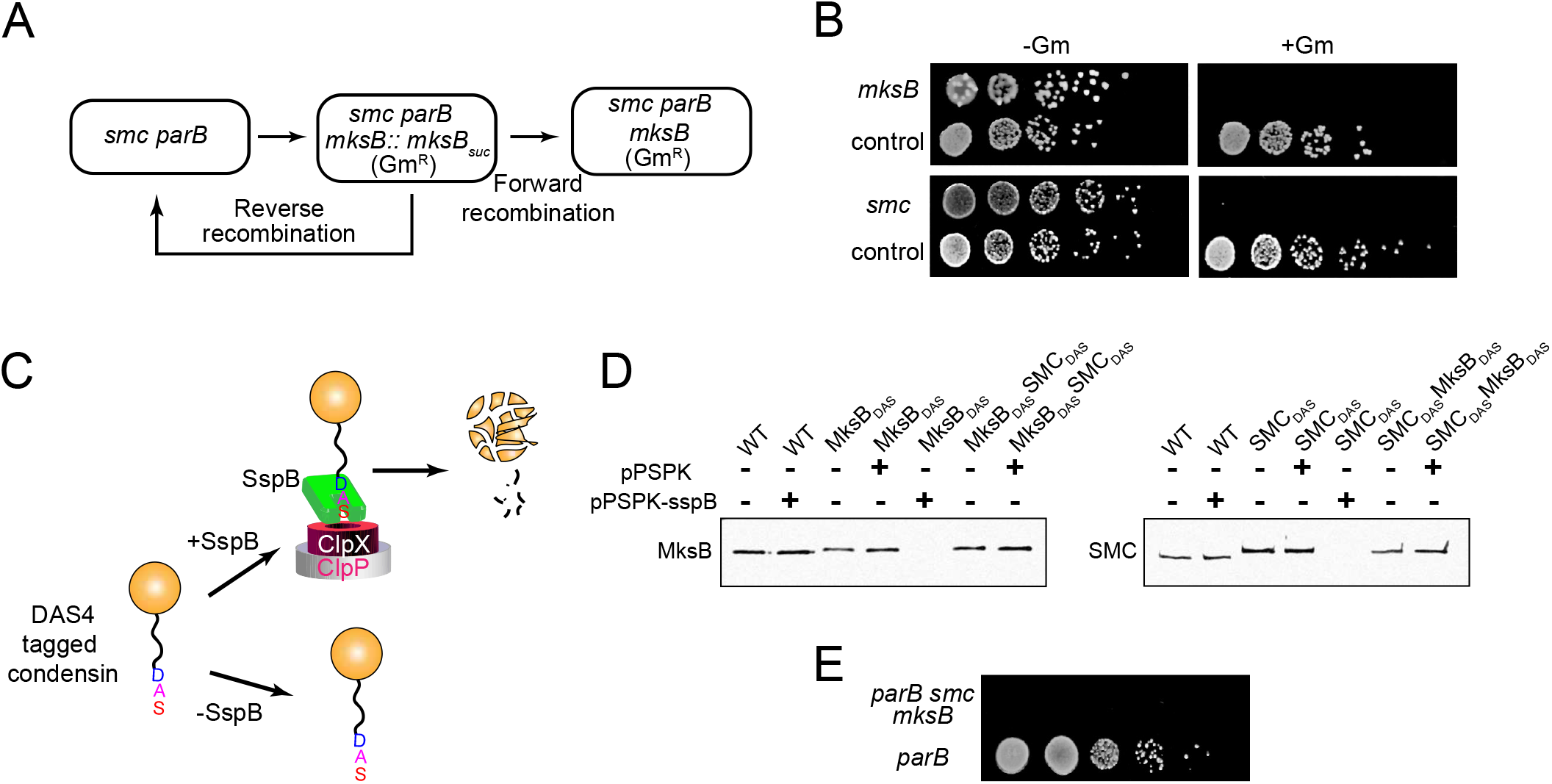
Condensins are synthetically lethal with ParB. (**A**) Two possible outcomes of a recombinational resolution of a condensin merodiploid strain. (**B**) Comparative analysis of the forward and reverse recombination in *mksB* (top panel) or *smc* (bottom panel) merodiploid strains. The strains were serially diluted and spotted onto LB plates supplemented with 15% sucrose and, when indicated, 15 μg/mL gentamicin. As a control, an unrelated PAO1 strain merodiploid in *mexGHID* operon, OP578, was used. (**C**) A diagram of controlled degradation of MksB-DAS4 and SMC-DAS4 proteins upon introduction of a SspB expression plasmid. (**D**) Immunoblotting analysis of SspB mediated degradation DAS4-tagged MksB and SMC in *ΔparB* cells. (**E**) SspB-mediated degradation of MksB and SMC in *ΔparB* cells is lethal. SspB expression plasmid was introduced into Δ*parB* Δ*sspB* cells harboring DAS4-tagged MksB and SMC. Cells were then spotted onto LB plates supplemented with gentamicin (30 μg/mL) and IPTG (0.1%). As a control, cells carrying an empty vector were also spotted onto the same plate.

To further validate this phenomenon, we employed a bacterial degron system (Fig. 6C), which involves degradation of specially tagged proteins by ClpXP protease (28, 29). To this end, the gene encoding an adaptor protein for the system, *sspB*, was removed from *ΔparB* cells, and the endogenous *mksB* and *smc* were replaced with their DAS4-tagged versions. In a permissive background, such replacement did not affect the expression levels of the condensins and resulted in a complete degradation of the proteins when a SspB-producing plasmid was introduced into the cells (Fig. 6D). When the plasmid was introduced into *ΔparB smc-das4 mksB-das4* cells, no colonies could be formed (Fig. 6E). For comparison, transformation of the cells with an empty vector did not impair their growth. We conclude, therefore, that inactivation of SMC, MksB and ParB is lethal for the cell. However, any one of the three systems is sufficient for cell viability.

## Discussion

Uniquely among model bacteria, *Pseudomonas aeruginosa* encodes two condensins, SMC and MksBEF. At the first glance, this arrangement is redundant. Even though SMC and MksBEF belong to two distinct superfamilies, their activities and functions are close enough to expect the same mechanism. We show here that this is not true. The two proteins have distinct localization patterns and their deletion produces discernable effects on chromosome structure. At the same time, each of the proteins can compensate for the absence of ParB, while at least one of them must be present to rescue ParB-deficient cells. Hence, MksBEF and SMC asymmetrically perform the same function in chromosome maintenance.

This function is dual in nature. On one hand, both SMC and MksBEF must be present to stabilize the macrodomain organization of PAO1-UW chromosome (Fig. 4). This feature implicates condensins in stabilization of the global chromosome organization. This conclusion is in accord with both biochemical and genomic studies. The ability of condensins to self-assemble into the chromosome scaffold has been deduced based on in vitro reconstitution studies.

Likewise, a Hi-C analysis of *E. coli* chromosome revealed the importance of condesins for a long range order within the chromosome (14). On the other hand, ParABS system compensated for the absence of both condensins, indicating that SMC and MksBEF also function in chromosome segregation (Fig. 6). Accordingly, the dynamics of chromosome segregation changed, again in an asymmetric manner, upon inactivation of SMC or MksB (Fig. 2E, 4). Thus, the activity of condensins simultaneously ensures global chromosome organization and segregation.

The fact that SMC and MksBEF perform this function in a distinct manner points to their parallel evolution, that the two proteins evolved separately to accomplish the same task. This conjecture naturally explains why some bacteria carry only SMC or MukBEF/MksBEF. Apparently, a single condensin suffices to ensure faithful chromosome segregation. At the same time, a complete exclusion of condensins from a synthetic genome adversely affected its stability, making condensins an obligatory member of a minimal bacterial genome. At the opposite extreme, many pathogenic and environmental bacteria encode multiple condensins.

Deletion of condensins from *P. aeruginosa* was detrimental for its pathogenicity (7). Clearly, having multiple condensins increases fitness of these bacteria in diverse environment.

Conversely, the loss of a condensin might help adapt bacteria to a specialized niche. For example, a spontaneous deletion of *mksE* and *mksF* from *P. aeruginosa* strain PAO1 created a subline PAO1-DSM, which also carries a chromosomal inversion (30) and is better adapted for a planktonic growth.

The coexistence of two condensins with partially redundant functions in *P. aeruginosa* helped reveal aspects of the proteins that were elusive in simpler systems. In particular, it became clear that each of the condensins contributes to the dynamics of *oriC* but in a unique manner. SMC was associated with the *oriC* region throughout the cell cycle except immediately after *oriC* duplication. One of the newly born *oriC*’s was essentially devoid of the condensin and became populated with it only with passage of time. Presumably, de novo synthesis of SMC or its recycling from the sister origin is needed to ensure full growth of the SMC cluster. MksB, on the other hand, remained away from *oriC* except shortly before its duplication. It is tempting to conclude from these data that the formation of MksB clusters at *oriC* is causatively linked to its duplication, either directly or indirectly. This could be because *oriC* replication creates a nucleation center for MksB, or vice versa, relocation of MksB triggers *oriC* duplication. Recently, a mechanism based on incompatibility of MukB clusters and *oriC* was proposed to explain chromosome segregation in *E. coli* (31).

Importantly, both SMC and MksB chased *oriC* in its relocation rather than led it. This eliminates the proteins as a plausible driving force for chromosome segregation. Consistent with this was the manner of condensin cluster formation at *oriC*. Both MksB and SMC clusters were highly dynamic and widely fluctuated around their centers. Their formation at new sites occurred abruptly, without clear intermediates or dissolution of clusters at the preceding location. Such dynamic is characteristic of systems involved in a reaction-diffusion patterning rather than translocation. It also befits a protein that modulates the activity of another rather than directly drags it around. Taken together, these data indicate that the role of condensins in chromosome segregation is more passive than active.

An important conclusion from this study is that condensins are not recruited to the *oriC* region, or the middle of the cell, or the cell quarters. They are recruited to all those locales, in a dynamic, growth phase dependent manner. This was especially clear for MksB, which was transiently recruited to *oriC* concurrent with its duplication and then remained at that site throughout the cell cycle and even past cell division. This dynamic marks cell quarters as a site where MksB clusters are born and the midcell as the place of their dispersal. In this respect, MksB serves as a chromosome organizing center that is born at the time and place of *oriC* duplication and remains there until the mother chromosome completes its replication.

Taken together, these data point to an elegant mechanism of chromosome segregation, which also could tie it to cell dynamics (Fig. 7). In this view, the life cycle of the cell has a beginning and an end. The cycle begins with duplication of the origins of replication, which quickly move apart via an as yet unknown mechanism. This duplication provides the initial driving force for chromosome segregation. The duplication is triggered by the convergence of MksB onto the site of replication, which marks the first special point inside the cell. In short cells with a single origin of replication, this special point lies in the middle of the cell. Cells with two *oriC*’s have two points of this type, which are located at the quarter positions, and so on. Further separation of *oriC*’s is probably driven by entropic forces, which take their roots in the increasing length of replicating chromosomes. Wherever *oriC*’s end up marks the second special point.

**Figure 7.**
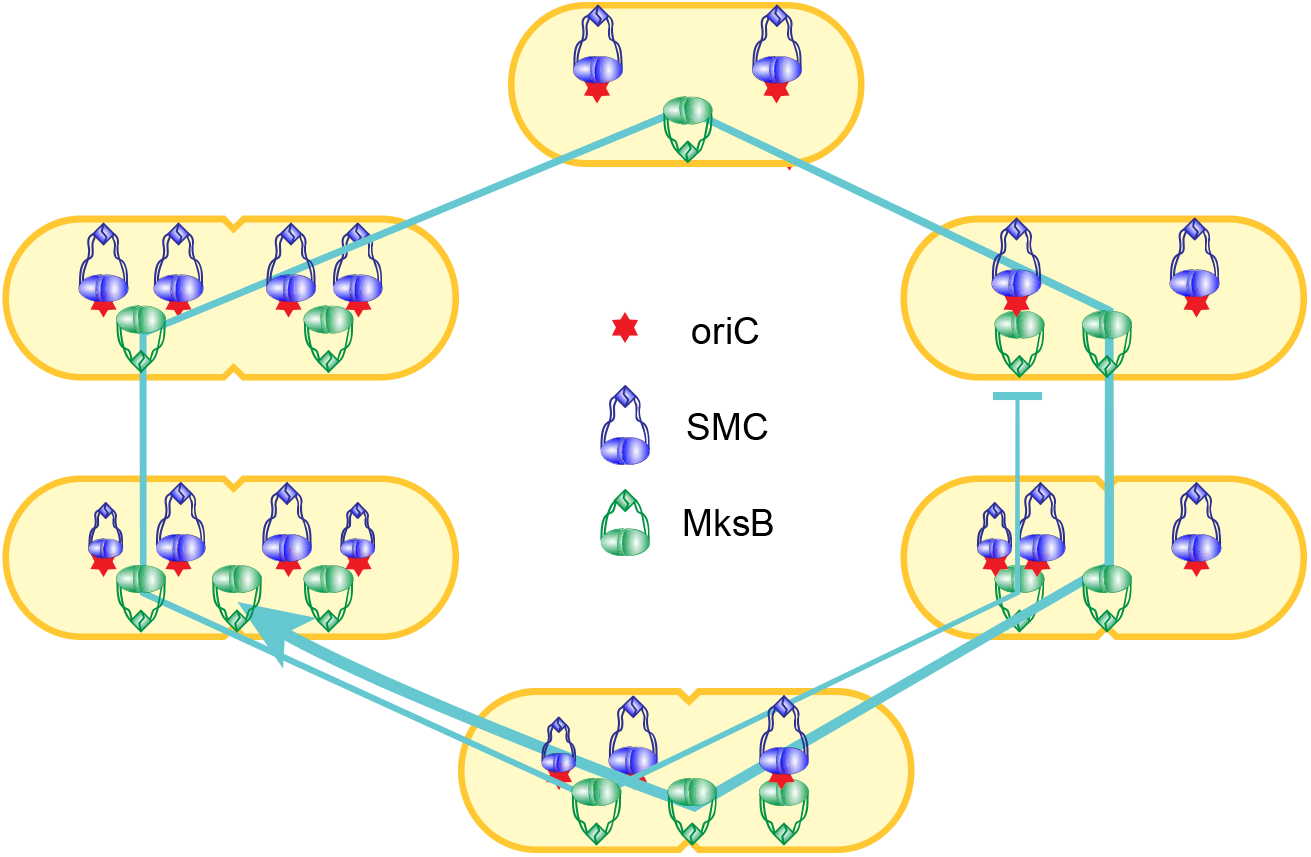
Dynamics of MksB, *oriC* and SMC during replication and segregation of the *P. aeruginosa* chromosome. The line traces the life cycle of MksB clusters, which form at the initiation of chromosome duplication and disperse upon completion of chromosome duplication and formation of both daughter duplication centers.

This second center is populated by SMC for the duration of the cell cycle, until MksB joins it. Thereafter, SMC and *oriC* leave it, and the site matures to become an organizing center for the daughter chromosome. It is quite possible that the daughter clusters coexist with the mother one for a period of time. In PAO1-UW cells in particular, the second round of *oriC* segregation occurs shortly before segregation of the *dif* region, which might explain the transient coexistence of three condensin clusters in one cell. The dispersion of the mother MksB cluster in the middle of the cell, which occurs after the daughters have already been born, marks the demise of the mother chromosome.

Perhaps the most intriguing result of this study is the found link between condensins and the cell architecture. Indeed, SMC was segregating with the septal rather than poleward *oriC*’s during the second round of *oriC* duplication (Fig. 5A). This pattern cannot occur by chance and depends on the SMC access to the information about the structure of the cell. Likewise, the deletion of any condensin resulted in a long delay in *oriC* segregation in one but not the other daughter cell (Fig. 2E). This asymmetry also implies structural differences in the cell caused by inactivation of SMC or MksB. The mechanism how condensins sense the architecture of the cell may offer clues to biogenesis of a living cell.

## Materials and Methods

### Plasmids and strains

*E. coli* strain DH5α (Novagen) was used as a host to construct all the plasmids, whereas *E. coli* strain SM10 (λ pir) was used for conjugation with *P. aeruginosa*. *P. aeruginosa* strains and plasmids used in this study are listed in Table S1 and S2. PAO1 (ATCC 47085) was used as the wild-type strain. Single and double condensin deletion mutant PAO1 strains were generated as described in (7). To tag the chromosomal loci in mutant strains, approximately 500 bp chromosomal segments at the desired locations were amplified using PCR and inserted between HindIII and KpnI restriction sites of the pP30DFRT-*tetO*-0069 plasmid. These plasmids carry approximately 140 *tetO* repeats. These plasmids were inserted into specific locations of the chromosome via homologous recombination.

Plasmid pPSV35Ap-TetR-mCherry was constructed by replacing the TetR-CFP from pPSV35Ap-TetR-CFP with its mCherry tagged version. This plasmid carries an in-frame fusion of TetR and mCherry genes under the control of isopropyl-β-D-thiogalactopyranoside (IPTG)-inducible *lacUV5* promoter. Chromosomal tags were visualized by introducing pPSV35Ap-TetR-CFP or pPSV35Ap-TetR-mCherry plasmids into the target cells via electroporation and selecting on LB agar plates supplemented with 200 μg/ml carbenicillin.

To construct pEX18Ap-SMC-mVenus plasmid, approximately 500 bp of the 3’-end of *smc* gene (PA1527) were linked to a construct encoding peptide HHHHHHHHHGG and *mVenus*. Then, this fused gene, the FRT-flanked gentamicin resistance cassette and 500 bp downstream of *smc* were inserted into the pEX18Ap plasmid. Similar approach was used to construct pEX18Ap-MksB-GFP and pEX18Ap-SMC-GFP plasmids. Endogenous copies of *mksB* and *smc* were replaced with their fluorescent tagged versions using allelic exchange method (32). Gentamicin resistance marker (*aac1*) was excised with the help of pFLP2 plasmid (33). Successful replacements were verified by PCR.

Plasmid pEXG2-ΔParB was constructed by amplifying approximately 500 bp chromosomal segments upstream and downstream of *parB* (PA5562). Resulting fragments were then cloned between Hind III and Pst I restriction sites of the shuttle vector pEXG2 (34). This plasmid was then used to create PAO1 Δ*parB* Δ*smc* (OP435), PAO1 Δ*parB* Δ*mksB* (OP441), PAO1 Δ*sspB* Δ*smc ΔparB* (OP475), PAO1 Δ*sspB* Δ*mksB ΔparB* (OP489) by allelic exchange. Deletions were subsequently confirmed by PCR. OP498 and OP497 was constructed by replacing the endogenous copies of *smc* and *mksB* in OP130, by their DAS4 tagged versions respectively. OP501 and OP508 were constructed by deleting *parB* from OP498 and OP132 respectively. OP505 and OP440 were constructed by deleting *mksB* and *smc* from OP501 and OP508 respectively. OP506 was constructed by replacing the endogenous copy of *mksB* gene by its DAS4 tagged version from OP501. OP529 and OP385 was constructed by introducing chromosomal tags at *oriC* proximal location following the method described above. PAO1 *ΔmexGHID* (*merodiploid*) (OP578) was constructed using allelic exchange method by mating PAO1 cells with SM10 cells carrying pEX18Ap-ΔGHID plasmid (35).

### Bacterial growth and microscopy

PAO1 cells harboring chromosomal tags and pPSV35Ap-TetR-CFP or pPSV35Ap-TetR-mCherry plasmids were grown overnight at 37°C in M9 minimal medium supplemented with 0.25% sodium citrate (M9+citrate) and a mixture of trace ions (36). Gentamicin (30 μg/ml) and carbenicillin (200 μg/ml) was added to the medium whenever necessary. Cells were then transferred to fresh M9+citrate medium at an OD_600_ of 0.01 and incubated at 30°C. After 3 hours of cell growth, expression of fluorescent-tagged TetR proteins was induced by adding 0.05 mM IPTG. Cells were collected at an OD600 of 0.1 and deposited on an agarose pad (1% agarose in M9+citrate medium) and promptly observed using a fluorescence microscope (Olympus BX50 equipped with a BX-FLA mercury light source, a 100X, 1.43 numerical-aperture oil immersion objective). Phase-contrast and fluorescent images were collected using a Spot Insight QE camera and analyzed using Nucleus and microbe (37, 38).

### Time-lapse imaging

For time-lapse imaging, cells were grown as described above. Cells were deposited onto agarose pads (1% agarose in M9+citrate medium) and snapshots were captured at 3-minute intervals using a fluorescence microscope (Olympus IX53 inverted microscope equipped with Sola light engine,). Images were captured using an ORCA-fusion digital camera (C14440).

Time-lapse image sequence was processed using FIJI (39) and home-written MATLAB software.

### Marker Frequency analysis

Marker frequency analysis was done as previously described (27). Briefly, the genomic DNA was isolated using a GenElute bacterial genomic DNA kit (Sigma-Aldrich). The DNA was then subjected to deep sequencing, and relative abundancies of each DNA site were binned and fitted to a model of bidirectional replication using a home-written MATLAB script. The rate of DNA replication normalized to cell growth rate (Mbp/μm) was determined from the fitting as one of the adjustable parameters of the model.

## Acknowledgments

We are indebted to Isabel Vallet-Gely for numerous insightful discussions and sharing unpublished data. We appreciate the Laboratory for Molecular Biology and Cytometry Research at OUHSC for performing the Illumina MiSeq sequencing and data analysis services for our whole-genome experiments. This work was supported in part by grants R21EY029015 and R21AI141927 from the National Institutes of Health to VVR.

## Author Contributions

H.Z. and B.B. performed the experiments. H.Z., B.B. and V.V.R analyzed data. V.V.R. wrote the manuscript and supervised the project.

## Declaration of Interests

The authors declare no competing interests.

## Supporting Information Legends

Supporting Information file includes Supplemental Figures S1 and S2 and Supplemental Tables S1and S2, which list strains and plasmids used in this study.

## Notes

### Competing Interest Statement

The authors have declared no competing interest.

## Reference

1. Wagner VE, Iglewski BH. 2008. P. aeruginosa Biofilms in CF Infection. Clin Rev Allergy Immunol 35:124–34.

2. Winstanley C, O’Brien S, Brockhurst MA. 2016. Pseudomonas aeruginosa Evolutionary Adaptation and Diversification in Cystic Fibrosis Chronic Lung Infections. Trends Microbiol 24:327–337.

3. Huang H, Shao X, Xie Y, Wang T, Zhang Y, Wang X, Deng X. 2019. An integrated genomic regulatory network of virulence-related transcriptional factors in Pseudomonas aeruginosa. Nat Commun 10:2931.

4. Balasubramanian D, Schneper L, Kumari H, Mathee K. 2013. A dynamic and intricate regulatory network determines Pseudomonas aeruginosa virulence. Nucleic acids research 41:1–20.

5. Hauser AR. 2011. Pseudomonas aeruginosa: so many virulence factors, so little time. Critical care medicine 39:2193–2194.

6. Zhao H, Clevenger AL, Ritchey JW, Zgurskaya HI, Rybenkov VV. 2016. Pseudomonas aeruginosa Condensins Support Opposite Differentiation States. J Bacteriol 198:2936–2944.

7. Rybenkov VV. 2014. Maintenance of chromosome structure in Pseudomonas aeruginosa. FEMS Microbiol Lett 356:154–65.

8. Nolivos S, Sherratt D. 2014. The bacterial chromosome: architecture and action of bacterial SMC and SMC-like complexes. FEMS Microbiol Rev 38:380–92.

9. Kleine Borgmann LA, Graumann PL. 2014. Structural maintenance of chromosome complex in bacteria. J Mol Microbiol Biotechnol 24:384–95.

10. Badrinarayanan A, Reyes-Lamothe R, Uphoff S, Leake MC, Sherratt DJ. 2012. In vivo architecture and action of bacterial structural maintenance of chromosome proteins. Science 338:528–31.

11. She W, Wang Q, Mordukhova EA, Rybenkov VV. 2007. MukEF is required for stable association of MukB with the chromosome. J Bacteriol 189:7062–8.

12. Kleine Borgmann LA, Ries J, Ewers H, Ulbrich MH, Graumann PL. 2013. The bacterial SMC complex displays two distinct modes of interaction with the chromosome. Cell reports 3:1483–92.

13. Lioy VS, Cournac A, Marbouty M, Duigou S, Mozziconacci J, Espeli O, Boccard F, Koszul R. 2018. Multiscale Structuring of the E. coli Chromosome by Nucleoid-Associated and Condensin Proteins. Cell 172:771–783 e18.

14. Surovtsev IV, Jacobs-Wagner C. 2018. Subcellular Organization: A Critical Feature of Bacterial Cell Replication. Cell 172:1271–1293.

15. Hiraga S, Ichinose C, Onogi T, Niki H, Yamazoe M. 2000. Bidirectional migration of SeqA-bound hemimethylated DNA clusters and pairing of oriC copies in Escherichia coli. Genes Cells 5:327–41.

16. Cobbe N, Heck MM. 2004. The evolution of SMC proteins: phylogenetic analysis and structural implications. Mol Biol Evol 21:332–47.

17. Petrushenko ZM, She W, Rybenkov VV. 2011. A new family of bacterial condensins. Mol Microbiol 81:881–96.

18. Wang X, Rudner DZ. 2014. Spatial organization of bacterial chromosomes. Curr Opin Microbiol 22:66–72.

19. Cui Y, Petrushenko ZM, Rybenkov VV. 2008. MukB acts as a macromolecular clamp in DNA condensation. Nat Struct Mol Biol 15:411–8.

20. Minnen A, Attaiech L, Thon M, Gruber S, Veening JW. 2011. SMC is recruited to oriC by ParB and promotes chromosome segregation in Streptococcus pneumoniae. Mol Microbiol 81:676–88.

21. Sullivan NL, Marquis KA, Rudner DZ. 2009. Recruitment of SMC by ParB-parS organizes the origin region and promotes efficient chromosome segregation. Cell 137:697–707

22. Nolivos S, Upton AL, Badrinarayanan A, Muller J, Zawadzka K, Wiktor J, Gill A, Arciszewska L, Nicolas E, Sherratt D. 2016. MatP regulates the coordinated action of topoisomerase IV and MukBEF in chromosome segregation. Nat Commun 7:10466.

23. Zawadzki P, Stracy M, Ginda K, Zawadzka K, Lesterlin C, Kapanidis AN, Sherratt DJ. 2015. The Localization and Action of Topoisomerase IV in Escherichia coli Chromosome Segregation Is Coordinated by the SMC Complex, MukBEF. Cell Rep 13:2587–2596.

24. Kumar R, Nurse P, Bahng S, Lee CM, Marians KJ. 2017. The MukB-topoisomerase IV interaction is required for proper chromosome compaction. J Biol Chem 292:16921–16932.

25. Vallet-Gely I, Boccard F. 2013. Chromosomal organization and segregation in Pseudomonas aeruginosa. PLoS genetics 9:e1003492.

26. Bhowmik BK, Clevenger AL, Zhao H, Rybenkov VV. 2018. Segregation but Not Replication of the Pseudomonas aeruginosa Chromosome Terminates at Dif. MBio 9.

27. McGinness KE, Baker TA, Sauer RT. 2006. Engineering controllable protein degradation. Mol Cell 22:701–7.

28. Castang S, McManus HR, Turner KH, Dove SL. 2008. H-NS family members function coordinately in an opportunistic pathogen. Proc Natl Acad Sci U S A 105:18947–52.

29. Klockgether J, Munder A, Neugebauer J, Davenport CF, Stanke F, Larbig KD, Heeb S, Schock U, Pohl TM, Wiehlmann L, Tummler B. 2010. Genome diversity of Pseudomonas aeruginosa PAO1 laboratory strains. J Bacteriol 192:1113–21.

30. Hofmann A, Makela J, Sherratt DJ, Heermann D, Murray SM. 2019. Self-organised segregation of bacterial chromosomal origins. Elife 8.

31. Hmelo LR, Borlee BR, Almblad H, Love ME, Randall TE, Tseng BS, Lin C, Irie Y, Storek KM, Yang JJ, Siehnel RJ, Howell PL, Singh PK, Tolker-Nielsen T, Parsek MR, Schweizer HP, Harrison JJ. 2015. Precision-engineering the Pseudomonas aeruginosa genome with two-step allelic exchange. Nature Protocols 10:1820.

32. Hoang TT, Karkhoff-Schweizer RR, Kutchma AJ, Schweizer HP. 1998. A broad-host-range Flp-FRT recombination system for site-specific excision of chromosomally-located DNA sequences: application for isolation of unmarked Pseudomonas aeruginosa mutants. Gene 212:77–86.

33. Rietsch A, Vallet-Gely I, Dove SL, Mekalanos JJ. 2005. ExsE, a secreted regulator of type III secretion genes in &lt;em&gt;Pseudomonas aeruginosa&lt;/em&gt. Proceedings of the National Academy of Sciences 102:8006.

34. Wolloscheck D, Krishnamoorthy G, Nguyen J, Zgurskaya HI. 2018. Kinetic Control of Quorum Sensing in Pseudomonas aeruginosa by Multidrug Efflux Pumps. ACS Infect Dis 4:185–195.

35. Schleheck D, Barraud N, Klebensberger J, Webb JS, McDougald D, Rice SA, Kjelleberg S. 2009. Pseudomonas aeruginosa PAO1 Preferentially Grows as Aggregates in Liquid Batch Cultures and Disperses upon Starvation. PLoS One 4:e5513.

36. Ducret A, Quardokus EM, Brun YV. 2016. MicrobeJ, a tool for high throughput bacterial cell detection and quantitative analysis. Nature Microbiology 1:16077.

37. Wang Q, Mordukhova EA, Edwards AL, Rybenkov VV. 2006. Chromosome condensation in the absence of the non-SMC subunits of MukBEF. J Bacteriol 188:4431–41.

38. Schindelin J, Arganda-Carreras I, Frise E, Kaynig V, Longair M, Pietzsch T, Preibisch S, Rueden C, Saalfeld S, Schmid B, Tinevez J-Y, White DJ, Hartenstein V, Eliceiri K, Tomancak P, Cardona A. 2012. Fiji: an opensource platform for biological-image analysis. Nature Methods 9:676.

