## supplemental figure 1 for "Alternating dynamics of *oriC*, SMC and MksBEF in segregation of *Pseudomonas aeruginosa* chromosome"

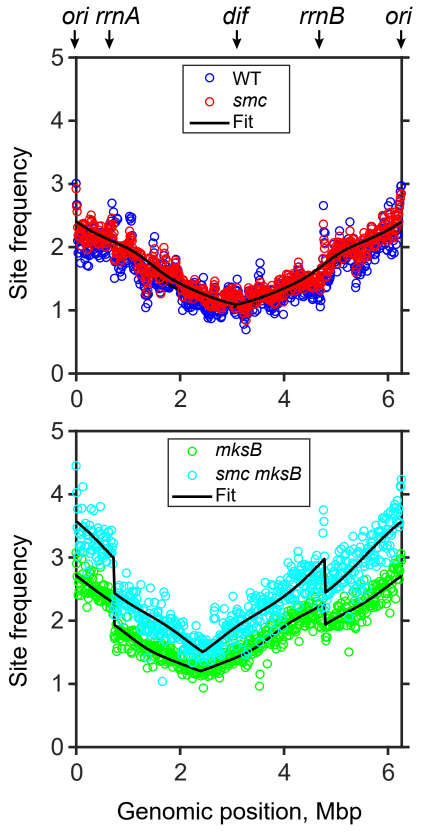


**Figure S1.** Marker frequency analysis of chromosome replication in condensin deficient cells without corrections for the chromosomal inversion. Genomic location of the quantified sites is shown using the chromosome of PAO1-UW as a reference. The discontinuities detected in *mksB* and *smc mksB* cells mark the junctions of the inversion.
