## supplemental figure 2 for "Alternating dynamics of *oriC*, SMC and MksBEF in segregation of *Pseudomonas aeruginosa* chromosome"

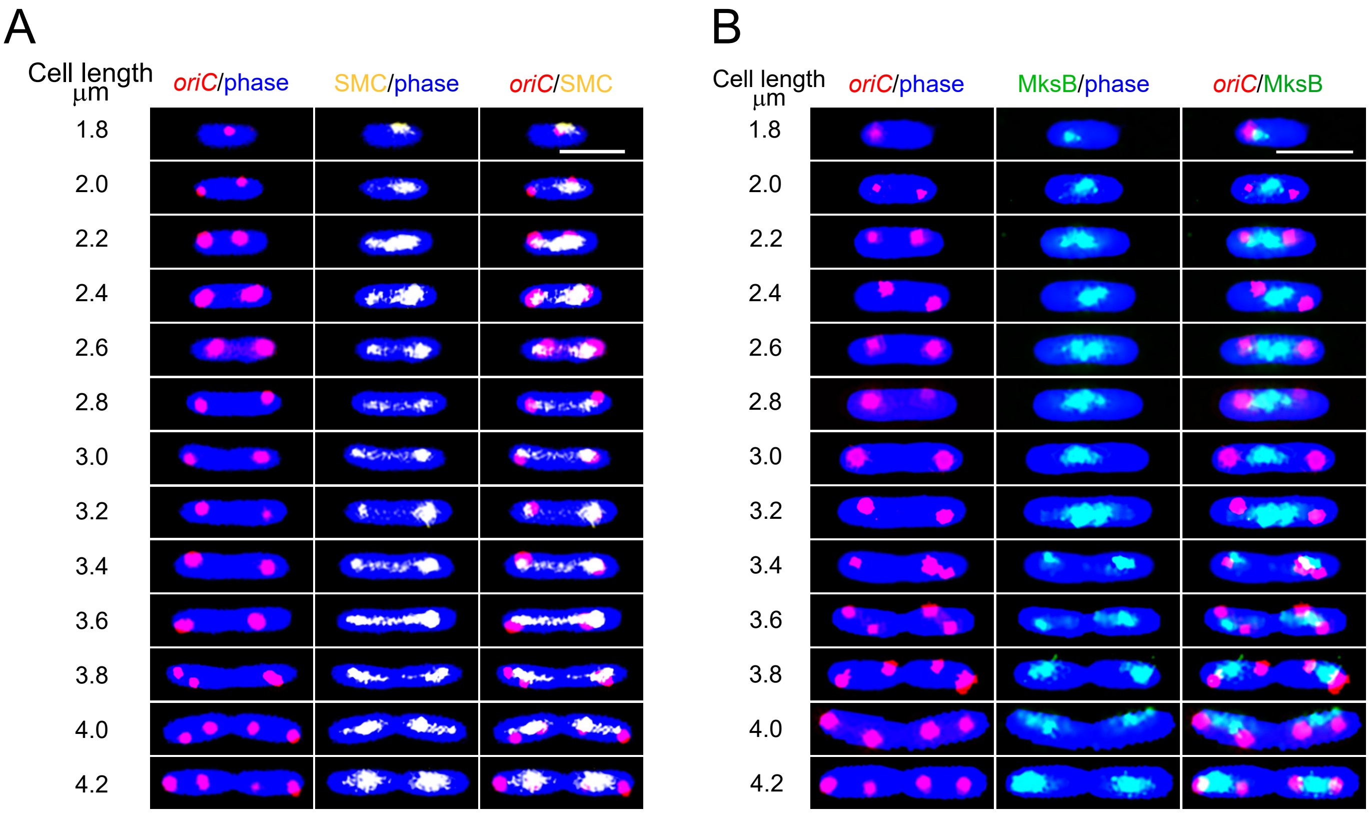


**Figure S2.** Representative cells with mCherry-tagged *oriC* and mVenus-tagged SMC (**A**) or mClover-tagged MksB (**B**). Scale bar, 2 µm.
