## supplemental Table 1 for "Alternating dynamics of *oriC*, SMC and MksBEF in segregation of *Pseudomonas aeruginosa* chromosome"

**Table S1.** Strains used in this study.

| Strain | Relevant genotype or description | Source or reference |
|---|---|---|
| DH5α | *supE44 ΔlacU169 hsdR17 recA1 endA1 gyrA96 thi-1 relA1* | Lab stock |
| SM10(λ pir) | *thi thr leu tonA lacY supE recA*::RP4-2-Tc::Mu Km λpir | Lab stock |
| PAO1-UW | *lacI*$^{q+}$ delta(*lacZ*)M15$^{+}$ *tetA*$^{+}$ *tetR*$^{+}$ | ATCC 47085 |
| PAO1-DSM | | (Klockgether et al., 2010) |
| OP107 | PAO1 Δ*smc* ΔGm | (Zhao et al., 2016) |
| OP109 | PAO1 Δ*mksB* ΔGm | (Zhao et al., 2016) |
| OP113 | PAO1 Δ*mksB* Δ*smc* ΔGm | (Zhao et al., 2016) |
| OP121 | PAO1 *mksB::mksB-GFP* Gm FRT | This study |
| OP123 | PAO1 *smc::smc-GFP* Gm FRT | This study |
| OP130 | PAO1 Δ*sspB* | (Zhao et al., 2016) |
| OP435 | PAO1 Δ*parB* Δ*smc* | This study |
| OP441 | PAO1 Δ*parB* Δ*mksB* | This study |
| OP475 | PAO1 Δ*sspB* Δ*smc* Δ*parB* | This study |
| OP489 | PAO1 Δ*sspB* Δ*mksB* Δ*parB* | This study |
| OP132 | PAO1 Δ*sspB mksB::mksB*-DAS4 ΔGm | (Zhao et al., 2016) |
| OP498 | PAO1 Δ*sspB smc::smc*-DAS4 ΔGm | This study |
| OP501 | PAO1 Δ*sspB* Δ*parB smc::smc*-DAS4 ΔGm | This study |
| OP508 | PAO1 Δ*sspB* Δ*parB mksB::mksB*-DAS4 ΔGm | This study |
| OP505 | PAO1 Δ*sspB* Δ*parB* Δ*mksB smc::smc*-DAS4 ΔGm | This study |
| OP440 | PAO1 Δ*sspB* Δ*parB* Δ*smc mksB::mksB*-DAS4 ΔGm | This study |
| OP506 | PAO1 Δ*sspB* Δ*parB smc::smc*-DAS4 *mksB::mksB*-DAS4 ΔGm | This study |

| | | |
|---|---|---|
| OP579 | PAO1 *smc::smc-mVenus* tetO-PA0069 + pPSV35Ap-TetR-mCherry | This study |
| OP385 | PAO1 *mksB::mksB*-GFP tetO-PA0069 + pPSV35Ap-TetR-mCherry | This study |
| OP461 | PAO1 tetO-PA0069 + pPSV35Ap-TetR-mCherry | This study |
| OP462 | PAO1 Δ*mksB* tetO-PA0069 + pPSV35Ap-TetR-mCherry | This study |
| OP457 | PA01 Δ*smc* PA0069 + pPSV35Ap-TetR-mCherry | This study |
| OP465 | PAO1 Δ*smc* Δ*mksB* tetO-PA0069 + pPSV35Ap-TetR-mCherry | This study |
| OP493 | PAO1 Δ*mksB* Δ*parB* Δ*smc (merodeploid)* | This study |
| OP487 | PAO1 Δ*smc* Δ*parB* Δ*mksB (merodeploid)* | This study |
| OP578 | PAO1 Δ*mexGHID (merodiploid)* | This study |
| BKB170 | PAO1 Δ*mksB* tetO-PA0069 + pPSV35Ap-TetR-CFP | This study |
| BKB243 | PAO1 Δ*mksB* tetO-PA0460 + pPSV35Ap-TetR-CFP | This study |
| BKB260 | PAO1 Δ*mksB* tetO-PA0716 + pPSV35Ap-TetR-CFP | This study |
| BKB175 | PAO1 Δ*mksB* tetO-PA0981 + pPSV35Ap-TetR-CFP | This study |
| BKB271 | PAO1 Δ*mksB* tetO-PA1436 + pPSV35Ap-TetR-CFP | This study |
| BKB286 | PAO1 Δ*mksB* tetO-PA1673 + pPSV35Ap-TetR-CFP | This study |
| BKB250 | PAO1 Δ*mksB* tetO-PA1905 + pPSV35Ap-TetR-CFP | This study |
| BKB174 | PAO1 Δ*mksB* tetO-PA2258 + pPSV35Ap-TetR-CFP | This study |
| BKB212 | PAO1 Δ*mksB* tetO-PA2910 + pPSV35Ap-TetR-CFP | This study |
| BKB287 | PAO1 Δ*mksB* tetO-PA3035 + pPSV35Ap-TetR-CFP | This study |
| BKB238 | PAO1 Δ*mksB* tetO-PA3267 + pPSV35Ap-TetR-CFP | This study |
| BKB171 | PAO1 Δ*mksB* tetO-PA3573 + pPSV35Ap-TetR-CFP | This study |
| BKB213 | PAO1 Δ*mksB* tetO-PA4457 + pPSV35Ap-TetR-CFP | This study |

| | | |
|---|---|---|
| BKB279 | PAO1 Δ*mksB* tetO-PA5099 + pPSV35Ap-TetR-CFP | This study |
| BKB326 | PAO1 Δ*smc* tetO-PA0069 + pPSV35Ap-TetR-mCherry | This study |
| BKB336 | PAO1 Δ*smc* tetO-PA0460 + pPSV35Ap-TetR-mCherry | This study |
| BKB328 | PAO1 Δ*smc* tetO-PA0716 + pPSV35Ap-TetR-mCherry | This study |
| BKB329 | PAO1 Δ*smc* tetO-PA0981 + pPSV35Ap-TetR-mCherry | This study |
| BKB337 | PAO1 Δ*smc* tetO-PA1436 + pPSV35Ap-TetR-mCherry | This study |
| BKB338 | PAO1 Δ*smc* tetO-PA1905 + pPSV35Ap-TetR-mCherry | This study |
| BKB315 | PAO1 Δ*smc* tetO-PA2258 + pPSV35Ap-TetR-mCherry | This study |
| BKB314 | PAO1 Δ*smc* tetO-PA2910 + pPSV35Ap-TetR-mCherry | This study |
| BKB320 | PAO1 Δ*smc* tetO-PA3267 + pPSV35Ap-TetR-mCherry | This study |
| BKB321 | PAO1 Δ*smc* tetO-PA3573 + pPSV35Ap-TetR-mCherry | This study |
| BKB312 | PAO1 Δ*smc* tetO-PA4457 + pPSV35Ap-TetR-mCherry | This study |
| BKB339 | PAO1 Δ*smc* tetO-PA5099 + pPSV35Ap-TetR-mCherry | This study |

### Supplemental References


Klockgether, J., Munder, A., Neugebauer, J., Davenport, C.F., Stanke, F., Larbig, K.D., Heeb, S., Schock, U., Pohl, T.M., Wiehlmann, L.*, et al.* (2010). Genome diversity of Pseudomonas aeruginosa PAO1 laboratory strains. J Bacteriol *192*, 1113-1121.

Zhao, H., Clevenger, A.L., Ritchey, J.W., Zgurskaya, H.I., and Rybenkov, V.V. (2016). Pseudomonas aeruginosa Condensins Support Opposite Differentiation States. J Bacteriol *198*, 2936-2944.
