## supplemental Table 2 for "Alternating dynamics of *oriC*, SMC and MksBEF in segregation of *Pseudomonas aeruginosa* chromosome"

**Table S2**. Plasmids used in this study.

| Plasmids | Description | Genomic location | Source or reference |
|---|---|---|---|
| pP30D-FRT-TetO-0069 | For insertion of *tetO* repeats at 12h (0.16h) | PA0069 | (Vallet-Gely and Boccard, 2013) |
| pP30D-FRT-TetO-0460 | For insertion of *tetO* repeats at 1h (0.99h) | PA0460 | (Bhowmik et al., 2018) |
| pP30D-FRT-TetO-0716 | For insertion of *tetO* repeats at 1.5h (1.51h) | PA0716 | (Bhowmik et al., 2018) |
| pP30D-FRT-TetO-0981 | For insertion of *tetO* repeats at 2h (2.04h) | PA0981 | (Vallet-Gely and Boccard, 2013) |
| pP30D-FRT-TetO-1436 | For insertion of *tetO* repeats at 3h (2.99h) | PA1436 | (Bhowmik et al., 2018) |
| pP30D-FRT-TetO-1673 | For insertion of *tetO* repeats at 3.5h (3.50) | PA1673 | This study |
| pP30D-FRT-TetO-1905 | For insertion of *tetO* repeats at 4h (3.98h) | PA1905 | (Bhowmik et al., 2018) |
| pP30D-FRT-TetO-2258 | For insertion of *tetO* repeats at 5h (4.76h) | PA2258 | (Vallet-Gely and Boccard, 2013) |
| pP30D-FRT-TetO-2910 | For insertion of *tetO* repeats at 6h (0.16h) | PA2910 | (Bhowmik et al., 2018) |
| pP30D-FRT-TetO-3035 | For insertion of *tetO* repeats at 6.5h (6.51h) | PA3035 | (Bhowmik et al., 2018) |
| pP30D-FRT-TetO-3267 | For insertion of *tetO* repeats at 7h (6.99h) | PA3267 | (Bhowmik et al., 2018) |
| pP30D-FRT-TetO-3573 | For insertion of *tetO* repeats at 8h (7.67h) | PA3573 | (Vallet-Gely and Boccard, 2013) |
| pP30D-FRT-TetO-4457 | For insertion of *tetO* repeats at 10h (9.6h) | PA4457 | (Bhowmik et al., 2018) |
| pP30D-FRT-TetO-5099 | For insertion of *tetO* repeats at 11h (10.99h) | PA5099 | (Bhowmik et al., 2018) |
| pPSV35Ap-TetR-CFP | Expressing TetR-CFP chimera | | This study |

| | | |
|---|---|---|
| pPSV35Ap-TetR-mCherry | Expressing TetR-mCherry chimera | This study |
| pFLP2 | Site-specific excision vector | (Hoang et al., 1998) |
| pEXG2 | deletion plasmid | (Castang et al., 2008) |
| pEXG2-Δ*sspB* | SspB deletion plasmid | (Castang et al., 2008) |
| pEXG2-Δ*parB* | ParB deletion plasmid | This study |
| pEX18AP | deletion plasmid | (Hoang et al., 1998) |
| pEX18Ap-Δ*smc* | SMC deletion plasmid | (Zhao et al., 2016) |
| pEX18Ap-Δ*mksB* | MksB deletion plasmid | (Zhao et al., 2016) |
| pEX18AP-*smc*-GFP | For chromosomal fusion protein | This study |
| pEX18AP-*mksB*-GFP | For chromosomal fusion protein | This study |
| pBAD-mVenus | Source of mVenus gene | Addgene |
| pEX18AP-*smc*-mVenus | For chromosomal fusion protein | This study |
| pEX18AP-*mksB*-DAS4 | ClpXP mediated Degradation of MksB protein | (Zhao et al., 2016) |
| pEX18AP-*smc*-DAS4 | ClpXP mediated Degradation of SMC protein | This study |
| pPSPK-*sspB* | IPTG inducible sspB expression plasmid | (Castang et al., 2008) |

### Supplemental References


Bhowmik, B.K., Clevenger, A.L., Zhao, H., and Rybenkov, V.V. (2018). Segregation but Not Replication of the Pseudomonas aeruginosa Chromosome Terminates at Dif. MBio *9*.

Castang, S., McManus, H.R., Turner, K.H., and Dove, S.L. (2008). H-NS family members function coordinately in an opportunistic pathogen. Proc Natl Acad Sci U S A *105*, 18947-18952.

Hoang, T.T., Karkhoff-Schweizer, R.R., Kutchma, A.J., and Schweizer, H.P. (1998). A broad-host-range Flp-FRT recombination system for site-specific excision of chromosomally-located

DNA sequences: application for isolation of unmarked Pseudomonas aeruginosa mutants. Gene *212*, 77-86.

Vallet-Gely, I., and Boccard, F. (2013). Chromosomal organization and segregation in Pseudomonas aeruginosa. PLoS genetics *9*, e1003492.

Zhao, H., Clevenger, A.L., Ritchey, J.W., Zgurskaya, H.I., and Rybenkov, V.V. (2016). Pseudomonas aeruginosa Condensins Support Opposite Differentiation States. J Bacteriol *198*, 2936-2944.
